# Early maternal loss affects diurnal cortisol slopes in immature but not mature wild chimpanzees

**DOI:** 10.1101/2020.10.15.340893

**Authors:** Cédric Girard-Buttoz, Patrick J. Tkaczynski, Liran Samuni, Pawel Fedurek, Cristina Gomes, Therese Löhrich, Virgile Manin, Anna Preis, Prince Valé, Tobias Deschner, Roman M. Wittig, Catherine Crockford

## Abstract

In mammals, early life adversity negatively affects survival and reproductive success. A key causal mechanism is proposed by the biological embedding model which posits that adversity experienced early in life has deleterious consequences on individual physiology across the lifespan. In particular, early life adversity is expected to be a severe stressor leading to long-term alteration of the hypothalamic pituitary adrenal (HPA) axis activity. Here we tested this idea by assessing whether, as in humans, maternal loss had short and long-term impacts on orphan chimpanzee urinary cortisol levels and diurnal urinary cortisol slopes, as an indicator of the HPA axis functioning. We used 18 years of data on 50 immature and 28 mature male wild chimpanzees belonging to four communities in Taï National Park, Ivory Coast. Immature orphans who experienced early maternal loss had diurnal cortisol slopes characterised by higher early morning and late afternoon cortisol levels indicative of high activation of the HPA axis. Recently orphaned immatures had higher cortisol levels than other immatures, possibly reflecting social and nutritional stress. However, unlike in humans, we did not find significantly different cortisol profiles in orphan and non-orphan adult male chimpanzees. Our study highlights that long-term alteration of stress physiology related to early life adversity may not be viable in some wild animal populations and/or that chimpanzees, as humans, may have access to mechanisms that buffer this physiological stress, such as adoption. Our results suggest that biological embedding of altered HPA axis function is unlikely to be a mechanism contributing to the demonstrated long-term fitness consequences of maternal loss, such as reduced reproductive success, in wild long-lived mammals.

## Introduction

In mammals, mothers are essential for the early development of their infants since they provide post-natal care (Maestripieri & Mateo 2009). Maternal loss in mammals reduces growth (Samuni et al. 2020), survival (Watts et al. 2009; Andres et al. 2013; Tung et al. 2016; Stanton et al. 2020), and long-term reproductive success (Andres et al. 2013; Strauss et al. 2020; Crockford et al. 2020, reviewed in Clutton-Brock 2016).

The biological embedding model (Power & Hertzman 1997; Miller et al. 2011; Berens et al. 2017) posits that adversity experienced early in life, including exposure to severe stressors, can have deleterious consequences on an individual’s physiology and health across their lifespan. This model provides a promising conceptual framework to investigate the mechanisms underlying the fitness costs of maternal loss or other forms of early life adversity.

Early life adversity impacts several interconnected physiological pathways (reviewed in Berens et al. 2017) among which the hypothalamic pituitary adrenal (HPA) axis plays a central role (Miller et al. 2009, 2011; Taylor et al. 2011). The HPA axis is activated in response to internal physiological challenges and external stressors through a chain of reactions, known as the “stress response” or “reactive homeostasis” (Romero et al. 2009), which also results in the release of glucocorticoids (Sapolsky 2002). Exposure to harsh social conditions during childhood, such as maternal loss, may lead to repeated and prolonged activation of the HPA axis early in life. These activations provide an adaptive physiological response by mobilising energy which helps children to cope with the immediate socio-ecological challenges but may result in long-term HPA axis dysfunction (i.e. either hypo- or hyper-responsiveness to stressor Miller et al. 2011; Ehrlich et al. 2016; Berens et al. 2017). The HPA axis is considered to be at the core of the link between early life adversity and fitness since repeated activation of the HPA axis over prolonged periods (chronic stress) and/or HPA axis malfunctioning can have detrimental effects on individual overall health (Sapolsky 2002; Slavich & Cole 2013). For instance, over- and/or prolonged activation of the HPA axis is known to suppress the immune system (Grossman 1985; Setchell et al. 2010; Slavich & Cole 2013) and the HPA axis mediates some of the observed negative effects of early life adversity on the immune response such as elevated levels of inflammatory markers in the blood (Danese et al. 2011; Ehrlich et al. 2016; Rasmussen et al. 2019, reviewed in Berens et al. 2017). Assessing the consequences of traumatic early life events, such as maternal loss, on the functioning of the HPA axis can provide insight into the mechanisms underlying the documented fitness costs of such events.

In humans, maternal loss leads to short and long-term alterations of the HPA axis functioning (Heim & Nemeroff 2001; Sánchez et al. 2001). These effects are typically studied by investigating patterns of cortisol excretion, the main glucocorticoid circulating in mammals, including humans. Cortisol levels follow a diurnal pattern characterized by an early morning peak (awakening response) and a regular decline throughout the day (Doman et al. 1986). In humans, diurnal cortisol slopes, and cortisol awakening response, serve as health markers. Deviations from the stereotypical patterns of high morning and low evening cortisol levels (i.e. flatter diurnal slopes) are typically interpreted as indications of pathology and HPA axis dysregulation and/or marker of chronic stress (Pruessner et al. 1999; Sánchez et al. 2001; Clow et al. 2004; Kudielka et al. 2006; Miller et al. 2007). Flattening of diurnal cortisol slopes reflect a compression in the dynamic range of the HPA axis functioning (Karlamangla et al. 2019) which is indicative of lowered ability to respond optimally to stressors and to down-regulate hormonal stress levels. In turn, flatter diurnal cortisol slopes may lead to direct fitness costs in humans such as reduced survival (Sephton et al. 2000).

To test conclusively the biological embedding model, the physiological consequences of maternal loss must be investigated during both during childhood and adulthood. Studies during childhood and in the time directly following maternal loss allow the assessment of the proximate responses of orphans in coping with new social challenges, while studies during adulthood allow the assessment of the long-lasting effect of early-life adversity. In humans, orphaned children typically exhibit lower cortisol awakening response (Carlson & Earls 1997) or higher evening cortisol levels (Gunnar et al. 2001) than mother-reared children, leading to overall flatter diurnal cortisol slopes (Carlson & Earls 1997; Tarullo & Gunnar 2006). Other forms of early life adversity, such as maltreatment by parents, parent divorce or placement in foster families, are also associated with flatter diurnal cortisol slopes (Kaufman 1991; Dozier et al. 2006; Bernard et al. 2015; McLachlan et al. 2016) and with lower morning cortisol levels (Hart et al. 1996; Meinlschmidt & Heim 2005). Early life adversity can also be associated with higher cortisol levels throughout the day, especially in children who experienced serious neglect (Cicchetti & Rogosch 2001; Gunnar et al. 2001; Wismer Fries et al. 2008).

In support of the biological embedding model, several studies in humans documented long-lasting effects of early life adversity on children’s HPA axis functioning (reviewed in Young et al. in press). Adults up to 64 years old who experienced mistreatment and/or the loss of one or both parents during childhood, depending on the study, have a lower (Meinlschmidt & Heim 2005; Kawai et al. 2017) or higher cortisol awakening response (Gonzalez et al. 2009; Butler et al. 2017), flatter diurnal cortisol slopes (Karlamangla et al. 2019), and generally higher cortisol levels throughout the day (Nicolson 2004).

Tests of the biological embedding model in wild long-lived animals with a slow life history are essential to understand if the long-lasting physiological effects of early life adversity found in humans are an artefact of our societies, which enable severely physiologically impaired individuals to survive until old age. Few studies have examined whether this impacted phenotypes are also present in adult wild mammals (Beehner & Bergman 2017) or whether HPA axis dysfunction simply leads to pre-adult death. Wild long-lived mammals are adapted to the environment in which we typically observe them, and in which selection may have favoured mechanisms of rapid recovery from early life traumatic events to avoid long-term hyper-activation of the HPA axis (or chronic stress, Beehner & Bergman 2017). In contrast, western humans, in which most studies on the impact of early life adversity on HPA axis functioning have been conducted, occupy a substantially modified environment compared to that in which humans evolved, including access to medical and institutional cares. As such, they may have reduced “recovery” ability due to lowered selective pressure associated with improved access to care.

A study on wild female baboons, using extensive long-term data, showed that simultaneous exposure to several forms of early life adversity, and some isolated forms of adversity such as drought and low maternal rank, leads to an overall elevation in glucocorticoid levels in adulthood (Rosenbaum et al. 2020), offering support for the biological embedding model. However, maternal loss in isolation did not lead to long-term elevation of glucocorticoid concentrations suggesting that baboons may have buffering mechanisms to offset effects of biological embedding for some forms of early life adversity. To our knowledge, this study on wild baboons constitutes the only test of the biological embedding model in a wild long-lived non-human mammal. More studies are necessary to investigate if, or how extensively, the biological embedding model applies to a wider range of long-lived species, both during development and in adulthood. In particular, it is important to test this model in long-lived species with a life history closer to that of humans. Baboons start reproducing one or two years after weaning, whereas humans and great apes, including chimpanzees share an extended juvenile phase between weaning and first reproduction (Wittig & Boesch 2019a). Furthermore, the study on baboons only assessed one marker of the HPA axis functioning (i.e. overall glucocorticoid levels) but did not investigate the impact on diurnal cortisol slopes. Assessment of diurnal cortisol slopes are important since these slopes are a marker of the HPA axis functioning (Karlamangla et al. 2019).

Using a long-term database, including demographic and urinary cortisol data, collected over a 20 year-period on four wild Western chimpanzee communities *(Pan troglodytes verus)* we aimed to provide rare test of the biological embedding model in a wild long-lived mammal. Specifically, our dataset allowed us to assess both the short and long-term effects of maternal loss on the HPA axis activity in wild chimpanzees. We thereby investigated one of the potential physiological mechanisms explaining the fitness costs associated with maternal loss reported in wild chimpanzees such as reduced growth, survival, and reproductive success (Nakamura et al. 2014; Samuni et al. 2020; Stanton et al. 2020; Crockford et al. 2020). Furthermore, studying physiological effects using diurnal cortisol slopes is an underused paradigm in wild animal subjects despite its prevalence in the human health literature. In chimpanzees, these slopes are repeatable in adults (i.e. are consistent within a given individual over time, Sonnweber et al. 2018) but also show plasticity to physiological challenges such as disease outbreaks (Behringer et al. 2020) or aging (Emery Thompson et al. 2020).

In humans, alloparental care such as placement in a foster family can reverse the physiological consequences of early life adversity (Gunnar et al. 2001). Like orphan humans, orphan chimpanzees may have access to buffering mechanisms such as alloparental care and support provided by conspecifics ranging from tolerance in feeding sites to full adoptions (i.e. daily consistent provisioning of cares to the orphans such as carrying, grooming, food sharing, Uehara & Nyundo 1983; Goodall 1986; Wroblewski 2008; Boesch et al. 2010; Hobaiter et al. 2014; Samuni et al. 2019). For immatures, we investigated the effect of maternal loss in both sexes. For adult individuals, we focused on males, the philopatric sex in chimpanzees (Pusey 1979; Boesch & Boesch-Achermann 2000), since the early life history of adult females who immigrated as adults into our study groups is often undocumented.

For both age-class groups (male and female immatures and adult males), we first assessed whether average cortisol levels and the steepness of the diurnal cortisol slopes differed between orphan and non-orphaned individuals. We predicted that, as in humans, immature orphans would exhibit higher overall cortisol levels and flatter diurnal cortisol slopes than non-orphans. We also predicted that, the overall effect of maternal loss on cortisol profiles would be more severe during the first years following maternal loss. That is because recently orphaned individuals have to adjust behaviourally and physiologically to a new social situation in which they do not benefit from maternal support and may have reduced access to food and socio-positive social interactions. Over time, orphans may adjust to, or may access compensatory strategies for, this new setup, which could result in lower impact on the HPA axis activity. Accordingly, following peak in alteration of the orphan cortisol profile directly after maternal loss, we anticipated some decline over time, but still for cortisol levels to remain elevated in orphans compared to non-orphans. We predicted this to last even into adulthood, matching the patterns of long lasting HPA activity alteration arising from maternal loss in humans and for other forms of early life adversity in wild baboons (Rosenbaum et al. 2020). Finally, we predicted that immature orphans that lost their mothers earlier in their lives would have flatter diurnal cortisol slopes and overall higher cortisol levels than immatures who lost their mother at a later age due to a greater level of dependency on mothers in early ontogeny (Clark 1977; Pusey 1983; Boesch & Boesch-Achermann 2000).

## Results

We used the long-term behavioural, demographic and urine sample data of the Taï Chimpanzee Project (Wittig & Boesch 2019b) collected on four communities of wild Western chimpanzees (East, North, Middle, and South) in the Taï National Park, Cote d’Ivoire (5°52’N, 7°20’E). The urine samples included in this study span over 19 years and were collected between 2000 and 2018. We used a series of Linear Mixed Models (LMMs) to test our predictions regarding the effect of maternal loss on overall cortisol levels and diurnal slopes (jointly constituting the cortisol profile) separately in immature males and females (i.e. individuals <12 years of age) and mature males (i.e. males > = 12 years of age). For both immatures and mature males, we tested first for differences in cortisol profiles between orphan and non-orphans (Model 1a for immatures and 1b for mature males) and, for orphans only, the effect of the age at which the orphan lost their mother on their cortisol profiles (Model 1b for immature orphans and 2b for mature male orphans). In addition, for immature orphans, we also tested the effect of the years since maternal loss (Model 1b).

In all the models, each urine sample represented a data point and the cortisol concentration of the sample (expressed in ng/ml SG) was the response variable. We used three predictor variables:

- *Orphan status:* Binary variable (yes/no) describing whether the immature was an orphan at the time of sample collection Model 1 a, and whether the adult male was orphaned before reaching 12 years of age (Model 2a).
- *Age when mother died:* Continuous variable describing the age at which immature orphans (Model 1b) and mature male orphans (Model 2b) lost their mother.
- *Years since maternal loss:* Continuous variable describing the number of years since immature orphans lost their mother (Model 1b)

For each of the models, all of these test predictors were included in interaction with both the linear and the quadratic terms for time of sample collection to test the effect of these test predictors on diurnal cortisol slopes. The quadratic term for time of sample collection was included here since a previous study showed that diurnal cortisol slopes in Taï chimpanzees follow a curved quadratic pattern (Sonnweber et al. 2018). In all our models we controlled for sex of the individual, community size, sex ratio of mature individuals in the community, age of the individual (except for Model 1b because of collinearity issue, see methods), the LCMS method used (“old” or “new” method, see the *urine analysis* section) and seasonal variation in ecological conditions (see methods). In addition, we controlled for repeated observations of the same individual over the same year by incorporating *individual ID* and *year* as random factors in each model. Finally, to control for the changes in cortisol diurnal slope with age, we built one slope per individual per year into each model by incorporating the dummy variable “individual_year” as a random factor.

### Effect of maternal loss on immature cortisol profiles

For Model 1a, assessing if immature orphans and non-orphans differ in their cortisol profiles, the full model was not significantly different from the null model (N = 849 samples and 50 individuals, LRT, df = 3, χ^2^ = 6.67, P = 0.083). However, since the p-value was close to the arbitrary threshold of 0.05 we investigated the significance of the predictors in the model, keeping in mind that the significance of these predictors should be interpreted with caution. In Model 1a, the significant interaction between *orphan status* and *time of the day* (LRT, P = 0.016, Table 2) suggests that immature orphans had a steeper daily cortisol slope than non-orphans (Figure 1). This difference in slopes may stem from higher early morning cortisol levels in immature orphans as compared to non-orphans (Figure 1). The marginal R^2^ and the conditional R^2^ for Model 1a were 0.259 and 0.606 respectively.

**Figure 1:**
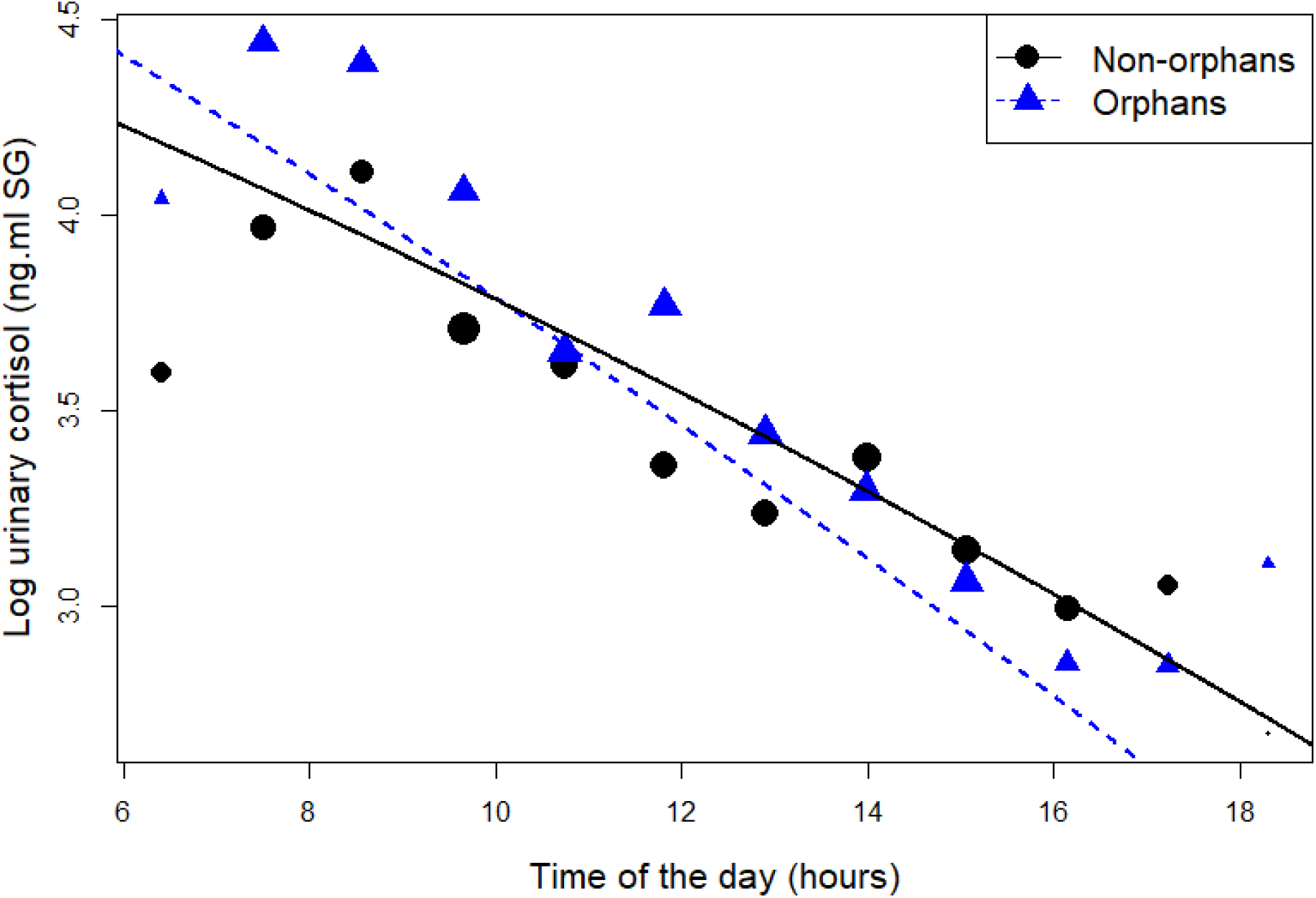
Effect of maternal loss on daily urinary cortisol level variations in immature chimpanzees. Orphans are depicted in blue triangles and non-orphans in black circles. The size of the dot is proportional to the sample size (e.g. the number of data points) contributing to each dot. The blue dotted line and black solid line depict the prediction lines from Model 1a for orphan and non-orphan respectively. Note that the full-null model comparison in Model 1a was not significant (P = 0.083) so the model lines should be interpreted with caution.

**Table 1:** Sample size for immature and adult male orphans and non-orphans in each of the 4 study communities.

| Community | Age class | Orphan status | N. individuals | N. samples | Age range (years) |
| --- | --- | --- | --- | --- | --- |
| Taï East | Immatures | non-orphans | 9* | 113 | 3.8 – 11.9 |
|  |  | orphans | 10* | 254 | 4.1 – 11.9 |
|  | Adult males | non-orphans | 4 | 456 | 13.8 – 40.7 |
|  |  | orphans | 3 | 354 | 12.3 – 19.8 |
| Taï Middle | Immatures | non-orphans |  |  |  |
|  |  | orphans |  |  |  |
|  | Adult males | non-orphans | 3 | 17 | 17.2 – 33.7 |
|  |  | orphans |  |  |  |
| Taï North | Immatures | non-orphans | 11 | 168 | 3.0 – 12.0 |
|  |  | orphans | 2 | 7 | 10.5 – 11.3 |
|  | Adult males | non-orphans | 3 | 95 | 12.3 – 20.8 |
|  |  | orphans | 2 | 61 | 12.1 – 20.4 |
| Taï South | Immatures | non-orphans | 17* | 173 | 2.8 – 11.9 |
|  |  | orphans | 5* | 134 | 4.1– 11.9 |
|  | Adult males | non-orphans | 7 | 858 | 12.1 – 45.3 |
|  |  | orphans | 6 | 343 | 12.1 – 21.9 |

\* 50 immature individuals were included in this study but two immatures in Taï East and two immatures in Taï South were sampled before and after their mother died (i.e. they are counted twice in the table, once as an orphan and once as a non-orphan). 6 males were included in the study as both mature and immature individuals.

**Table 2:**
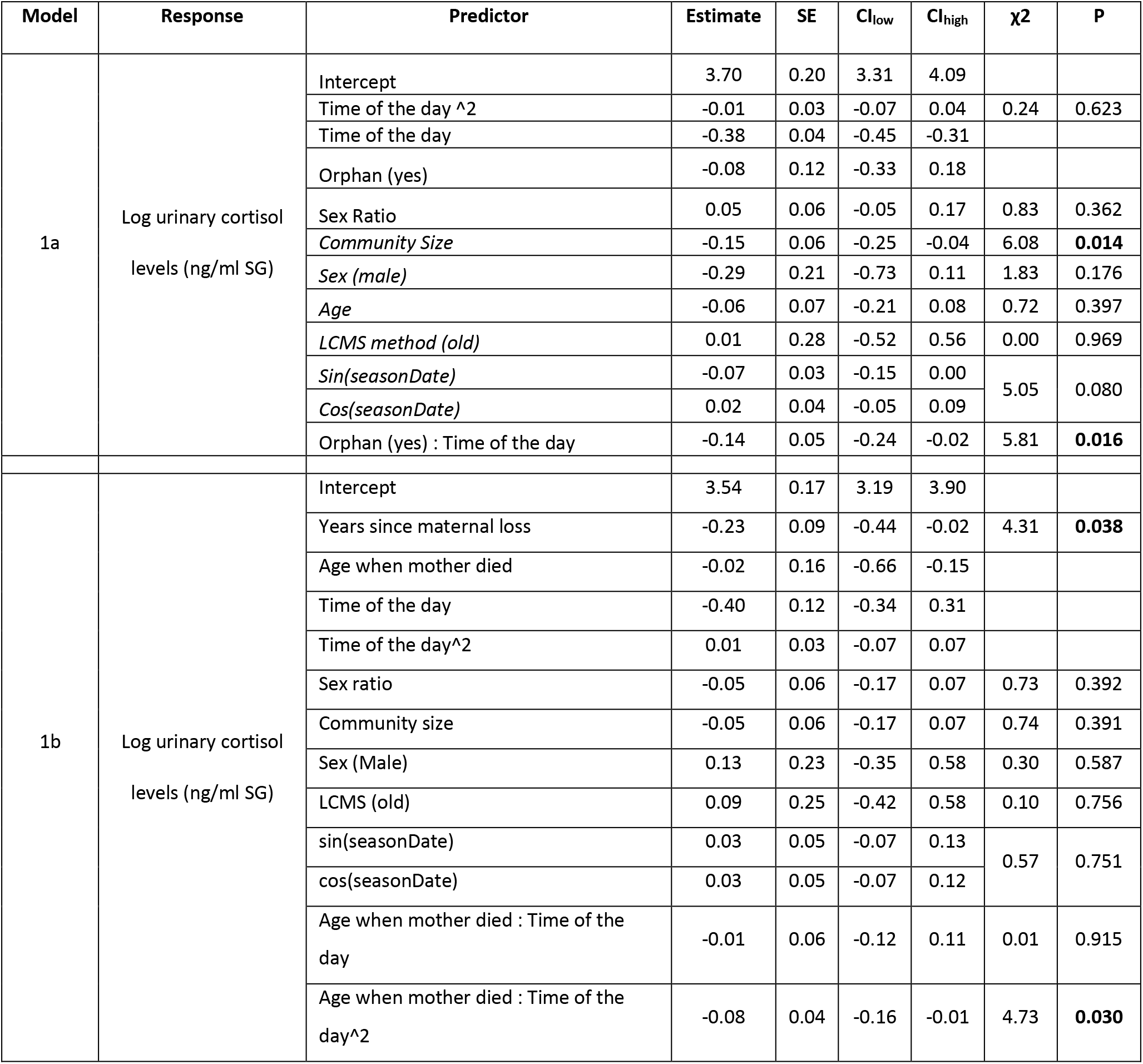
Results of Model 1a and 1 b. SE indicates the standard error of the estimate for each predictor. The coded level for each categorical predictor is indicated in parentheses. Control predictors are italicized. Significant p-value (p < 0.05) are indicated in bold. Ch_low_ and CI_high_ indicate the lower and upper limits of the 95% confidence interval for the estimates of each predictor. *For Model 1a the full-null model* comparison was only a trend (χ^2^ = 6.67, df = 3, P = 0.083) but since *it was close to the arbitrary significant level of 0.05 we nevertheless present the results of the single predictors in the table. The detailed results of model 2a and 2b are not presented since the full-null model comparison was not significant (both P> 0.4).*

The second model (Model 1b) focusing on immature orphans revealed that orphans vary significantly in their cortisol profiles depending on the age at which their mother died and on the length of time since they were orphaned (full-null model comparison in Model 1b, N = 393 samples, and 17 individuals, LRT, df = 6, χ^2^ = 12.82, P = 0.046). More specifically, immature orphans whose mother had died several years before sampling had significantly lower cortisol levels than more recently orphaned immatures (effect of *years since maternal loss* in Model 1b, β±SE = −0.23±0.09, P = 0.038, Table 2, Figure 2). To visualise this effect, we computed the average cortisol levels of individuals orphaned for less than 1 year, 1-2 years, 2-3 years, and 3-4 years as compared to the cortisol levels of aged-matched non-orphaned chimpanzees (Figure 3). The effect of *orphan status* on cortisol levels was apparent for individuals orphaned for less than two years at the time of sample collection, but orphans returned to “non-orphan” cortisol levels between two and three years after maternal loss (Figure 3).

**Figure 2:**
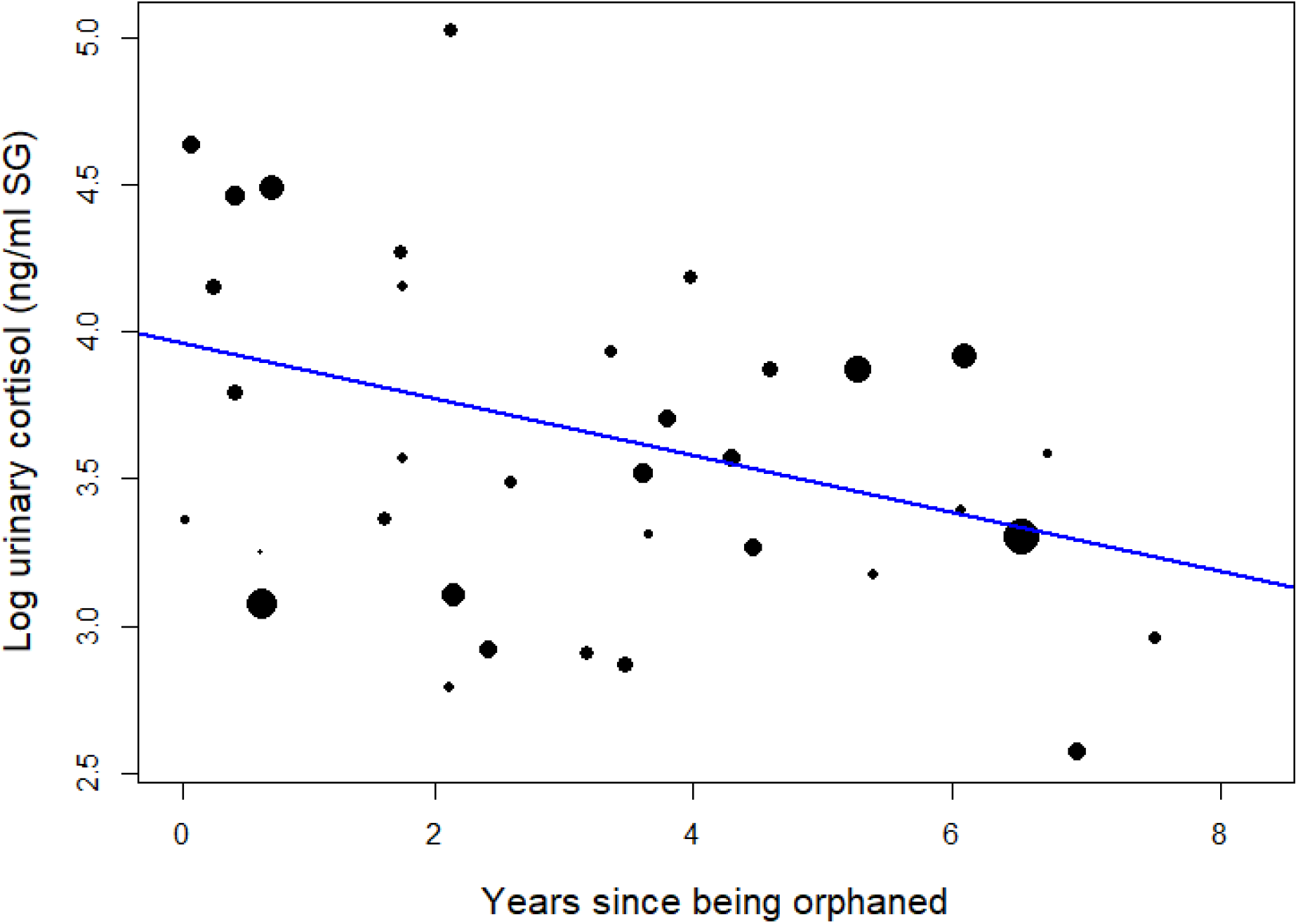
Effect of time (in years) since being orphaned on the urinary cortisol levels of immature orphan chimpanzees. Each dot represents each individual each calendar year it has been sampled. The size of the dot is proportional to the sample size (number of urine sample collected) for that individual that year. The blue line indicates the model line (Model 1b).

**Figure 3:**
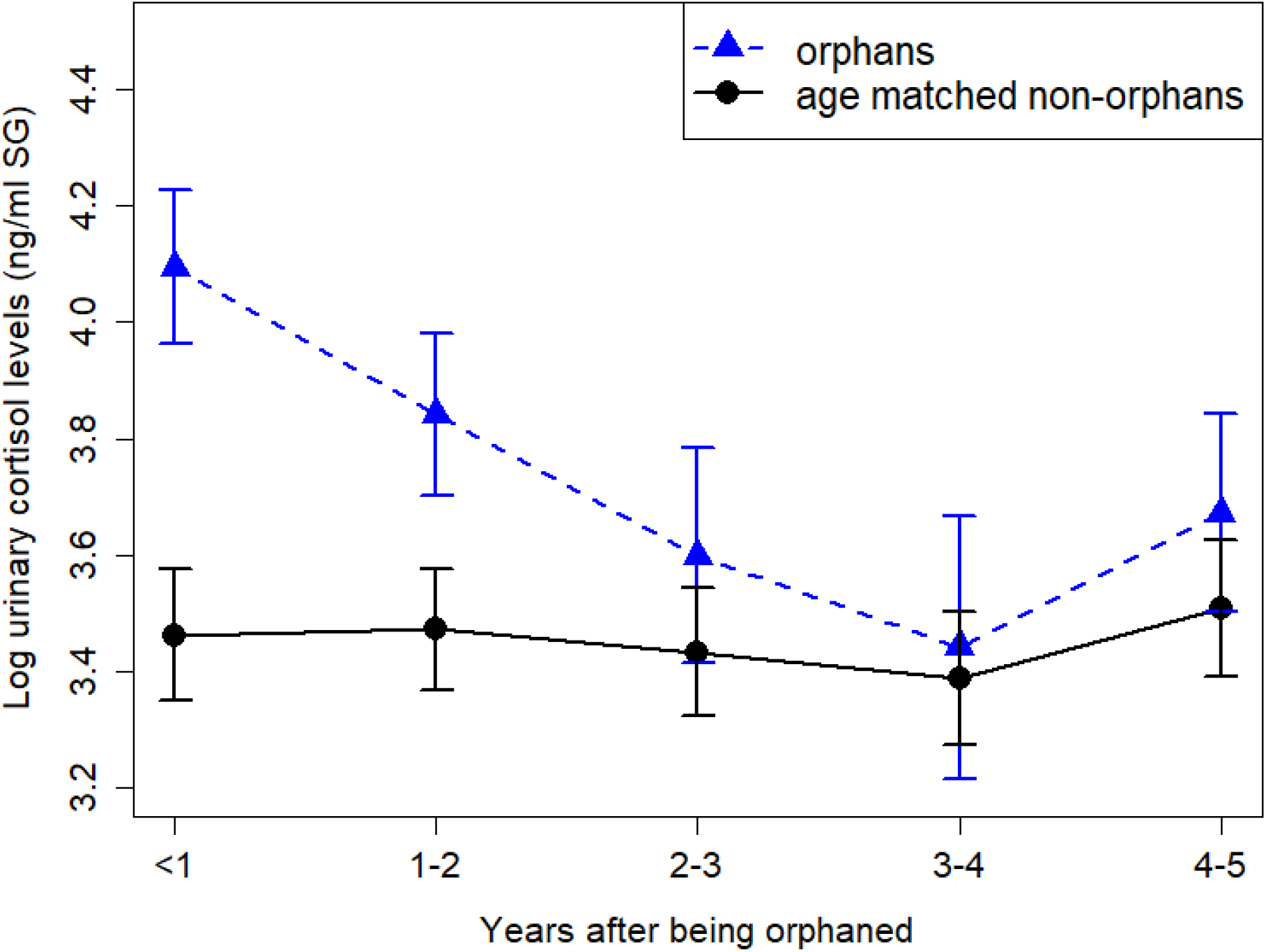
Effect of time (in years) since being orphaned on the urinary cortisol levels of immature orphan chimpanzees compared to age-matched non-orphan immatures. The mean ± se is depicted for orphans in blue triangles and for non-orphan in black circles. The non-orphan pattern indicates the mean urinary cortisol of non-orphan immature who fell into the age range of orphan immature who have been orphaned for less than a year, 1-2 years, 2-3 years, 3-4 years and 4-5 years respectively.

In addition, in Model 1b, significant interaction between *age when mother died* and quadratic term of *time of day* (β±SE = −0.08±0.04, P = 0.030, Table 2) indicates that the age at which immatures lost their mother significantly influenced their diurnal cortisol slopes. Immature individuals who lost their mother before the age of 5 years old (in orange Figure 4) had a diurnal cortisol slope that curved upwards and presented higher early morning and late afternoon cortisol levels than individuals who lost their mother at an older age. Those who lost their mother between 5 and 8 years of age (in blue in Figure 4) had a relatively linear decrease of cortisol throughout the day.

**Figure 4:**
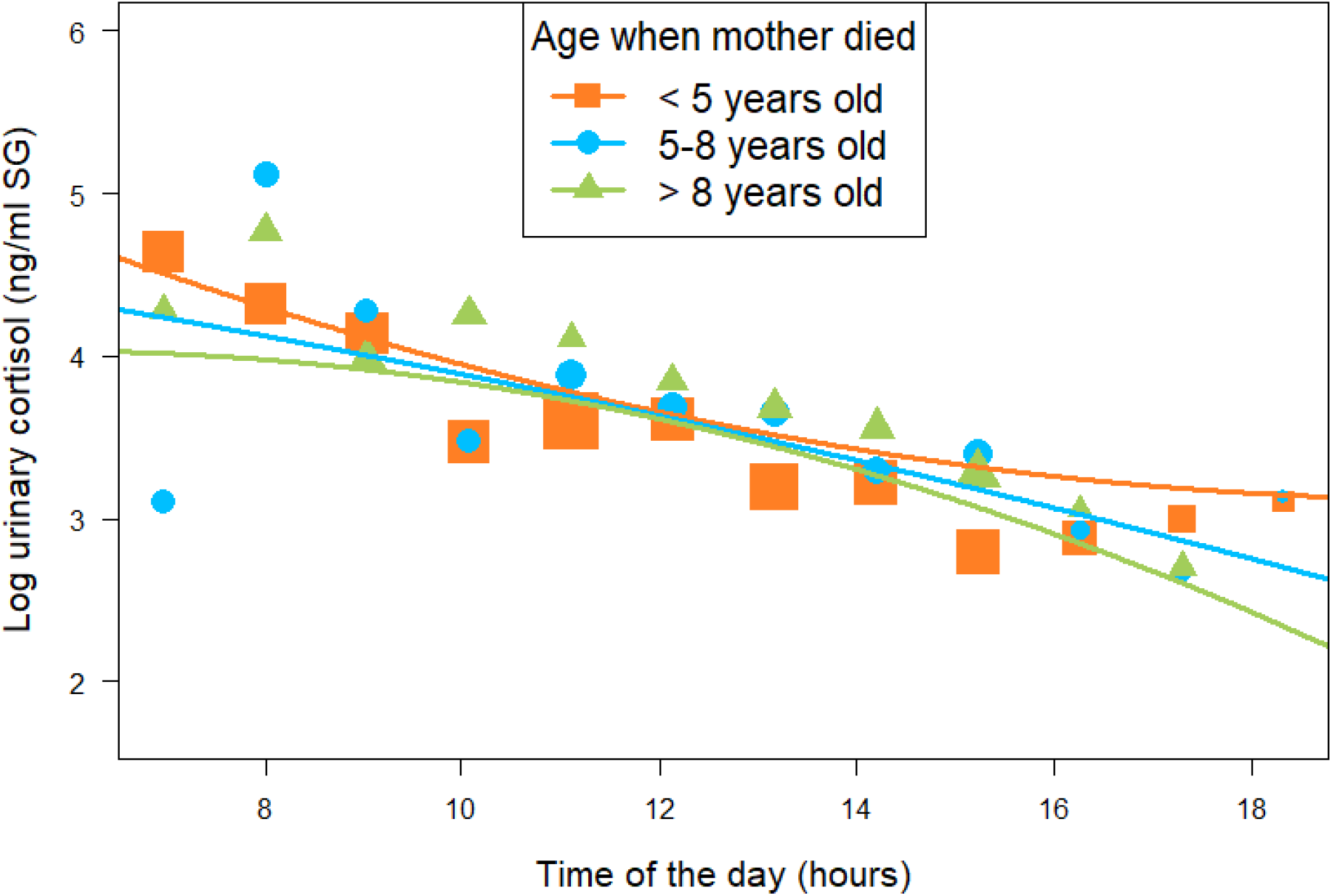
Effect of age at which immature orphans lost their mother on diurnal cortisol slopes. Each dot represents each individual each calendar year it has been sampled. The lines represent the model line predictions (Model 1b). The data points and model line prediction are depicted for immatures who have been orphaned before they were 5 years of age in orange squares, for immatures who have been orphan when they were between 5 and 8 years of age in blue circles, and for immatures who have been orphaned when they were older than 8 years of age in green triangles.

Finally, immature individuals orphaned after 8 years of age (in green in Figure 4) had a diurnal cortisol slope that curved downwards with lower early morning and late afternoon cortisol levels than individuals who lost their mother at a younger age. Please note that we depicted model line predictions for three age categories corresponding to three life history stages in chimpanzees (0-5 years: infancy, 5-8 years: juvenile period, 8-12 years: early adolescence) in Figure 4 for ease of interpretation of the effect but the variable “age when mother died” was incorporated in Model 1b as a continuous predictor. The marginal R^2^ and the conditional R^2^ for Model 1b were 0.235 and 0.594 respectively.

The results of Model 1b could potentially be explained via an effect of age on cortisol profiles in immatures. However, the *age of the individual* could not be incorporated as control factor in Model 1b due to collinearity with *age when mother died* and *“years since maternal loss”.* Therefore, we ran a separate model (Model 1c) to investigate the effect of age on cortisol levels and diurnal cortisol slopes in non-orphan immatures. In this model the full model was not significantly different from the null model (N = 454 samples, and 37 individuals, LRT, df = 3, χ^2^ = 1.98, P = 0.576). This indicates that the age of an immature did not significantly influence cortisol levels (as previously shown in the same population, Tkaczynski et al. 2020) and diurnal cortisol slopes in our study samples.

### Effect of maternal loss on cortisol slopes in mature male chimpanzees

In contrast to immature individuals, we did not detect a significant overall effect of *orphan status* on mature males’ cortisol profiles (full-null model comparison in Model 2a, N = 2184 samples, and 28 individuals, LRT, df = 3, χ^2^ = 0.51, P = 0.917). We also did not detect a significant effect of the *age when mother died* on orphan mature males’ cortisol profiled (full-null model comparison in Model 2b, N = 769 samples, and 10 individuals, LRT, df = 3, χ^2^ = 2.43, P = 0.488).

## Discussion

While the effect of maternal loss on wild animal survival and reproduction has been recently established (Foster et al. 2012; Andres et al. 2013; Tung et al. 2016; Walker et al. 2018; Surbeck et al. 2019; Crockford et al. 2020), the mechanisms underlying these fitness costs remain understudied. Our study provides one of the rare empirical tests of the biological embedding model in wild long-lived mammals (see also Rosenbaum et al. 2020), by assessing the short- and longterm physiological impacts of early maternal loss. While we found an effect of maternal loss on both overall cortisol levels and diurnal cortisol slopes in immature chimpanzees, these effects were not present in mature male chimpanzees. These results are in line with the absence of long-term effects of maternal loss alone on glucocorticoid levels in wild long-lived baboons (Rosenbaum et al. 2020). This suggests that 1) the biological embedding model (Power & Hertzman 1997; Miller et al. 2011; Berens et al. 2017) may apply to long-lived wild mammals only following exposure to a combination of diverse sources of early life adversity or to other sources of early life adversity than maternal loss (see Rosenbaum et al. 2020) and/or 2) that buffering mechanisms ranging from minimal alloparental cares to adoption (discussed below) could ameliorate long-term effects of early life adversity in chimpanzees. Alternatively, in our study, survivorship bias may mean that chimpanzees with severely altered HPA axis activity following maternal loss did not survive to adulthood and thus were absent from our adult dataset.

In contrast to our prediction, the difference between immature orphan and non-orphan diurnal cortisol slopes were marginal. Furthermore, immature orphans had a steeper diurnal cortisol slope than immature non-orphans, and this effect appeared to be driven principally by elevated early morning cortisol levels in orphans (Figure 1). This contrasts with findings in human studies that show flatter slopes for individuals that experience adversity (Kaufman 1991; Hart et al. 1996; Meinlschmidt & Heim 2005; Dozier et al. 2006; Bernard et al. 2015; McLachlan et al. 2016). A large portion of the variance in Model 1a (36%) was explained by the random effects indicating that the lack of significance in this analysis may be related to large inter-individual differences, and in particular largely varying cortisol levels and diurnal cortisol slopes across immature orphans. This is possibly due to buffering strategies available to some but not all orphans (e.g. whether the orphan was adopted or not) and to variation in the exposure to energetic and/or psychological stressors for different orphans due to several factors such as the age of the orphan when its mother died. In fact, orphans that lost their mother at a younger age had diurnal cortisol slopes characterised by higher early morning and late afternoon cortisol levels when compared to immatures orphaned when they were older. We also found a strong elevation in average cortisol levels in the year directly following maternal loss, followed by a return to cortisol levels comparable to age-matched non-orphans within 2-3 years after maternal loss.

Our results indicate that immature chimpanzees who lost their mother at a young age and/or recently lost their mothers experience unusually high levels of environmental and social stressors that challenge their homeostasis (Romero et al. 2009). Most likely, these stressors are social and energetic in nature, reflecting the lack of care and access to food sources that was initially provided by the mother (Pusey 1983; Goldenberg & Wittemyer 2017; Samuni et al. 2019). Even after weaning (which is the case in our study since all orphans were sampled after the weaning age c.a. 4 years of age, Samuni et al. 2020, Table S1), orphans may be constrained in acquiring the amount of food necessary to maintain a positive energy balance as they also lack food sharing and socially facilitated access to high nutrient food sources (Samuni et al. 2019). The elevated early morning cortisol levels found in orphans who lost their mother at a younger ages could be a sign of dietary restrictions (Goodwin et al. 1988; Garcia-Belenguer et al. 1993). In the same population, orphans lose out on growth compared to non-orphans (Samuni et al. 2020). Dietary restriction may thus also explain why immature orphan chimpanzees tended to have steeper diurnal cortisol slopes and higher early morning levels than non-orphans.

In contrast, in humans, orphans or infants who experienced other forms of early life adversity typically have blunted diurnal cortisol slopes and lower awakening morning responses (Kaufman 1991; Hart et al. 1996; Meinlschmidt & Heim 2005; Dozier et al. 2006; Bernard et al. 2015; McLachlan et al. 2016). This difference between chimpanzees and humans may stem from higher exposure to nutritional stress in immature orphan chimpanzees, as compared to orphan humans (at least in the Western human populations studied). Nutritional stress may also explain why we found that immatures who lost their mothers at a younger age had significantly higher afternoon cortisol levels and that the average cortisol levels (i.e. throughout the day) of orphans were elevated during the first two years following maternal loss.

These higher afternoon and average cortisol levels may also reflect exposure of orphans to social stress. In fact, immature chimpanzees who have lost their mother play for shorter periods of time (Botero et al. 2013), and play bouts escalate more frequently into aggression (van Leeuwen et al. 2014). This could, in turn, lead to increased cortisol levels since in chimpanzees and other primates, aggression generally increases cortisol levels (Girard-Buttoz et al. 2009; Emery Thompson et al. 2010; Wittig et al. 2015; but see Preis et al. 2019). Furthermore, strong social relationships, which some orphans may lack, can buffer the effect of environmental stressors on cortisol secretion in primates (Young et al. 2014; Wittig et al. 2016). Cumulatively, this suggests that new orphans and/or those who lost their mother at a young age may be more exposed to social and psychological stressors as well as reduced access to social mechanisms to buffer such stressors. The social factors affecting chimpanzee orphans may be similar to those impacting human orphans who are frequent targets of assault, either physical or sexual due to a lack of social support from a caregiver (e.g. Frank et al. 1996). It is thus not surprising that, overall, our results are in line with human and captive rodent studies showing that early life adversity elevates immature glucocorticoid levels (Cicchetti & Rogosch 2001; Gunnar et al. 2001; Meaney & Szyf 2005; Wismer Fries et al. 2008; Zhang et al. 2013).

While the effects of maternal loss on immatures physiology are comparable in humans and chimpanzees, the chimpanzee pattern differs strongly from that of humans, in that the effect of maternal loss on cortisol secretion profile did not persist into adulthood. Adult humans up to 64 years old, that experienced mistreatment and/or the loss of one or both parents during childhood, still present alteration of their diurnal cortisol slopes and overall cortisol levels (Nicolson 2004; Meinlschmidt & Heim 2005; Gonzalez et al. 2009; Kawai et al. 2017; Butler et al. 2017; Karlamangla et al. 2019). The re-establishment of a normal functioning of the HPA axis in mature male chimpanzees may reflect a form of recovery in those males. However, in our sample of mature males, we could, by definition, only sample individuals who survived until maturity. As a result, we cannot rule out the possibility that the orphaned infants most severely affected by maternal loss and who faced the strongest social and nutritional stress died at a young age and never reached adulthood. Unfortunately, we were only able to sample two immature chimpanzees the year before they died so we cannot evaluate the effect of cortisol profiles on individual survival.

Our study showed that individuals who lost their mother at a younger age had the most altered functioning of their HPA axis which could be one of the mechanisms explaining why other studies found that the younger an individual is when orphaned the less likely it is to survive into adulthood (Nakamura et al. 2014; Stanton et al. 2020). Furthermore, long-term alteration of the HPA axis functioning related to early life adversity is explained, at least partly, by epigenetic mechanisms (Weaver et al. 2004) and in particular by a hyper methylation of DNA in regions coding for the glucocorticoid receptors (GR) in the brain (Liu et al. 1997; Weaver et al. 2007, reviewed in Zhang et al. 2013). In rodents tested in laboratory conditions, these alterations of the GR in the brain take place early in life; a lower density of GR reduces the effectiveness of the glucocorticoid negative feedback loop and ultimately results in prolonged elevated cortisol levels (Zhang et al. 2013). All orphan adult males in our study were orphaned after they were four years of age, an age at which the epigenetic effect on GR in the brain may be reduced or absent. In addition, in captive rodents, this effect can be reversed following cross-fostering (Weaver et al. 2004),which may be equivalent to alloparental care or adoption in wild chimpanzees.

In chimpanzees, adoption is a common phenomenon (Uehara & Nyundo 1983; Goodall 1986; Wroblewski 2008; Boesch et al. 2010; Hobaiter et al. 2014; Samuni et al. 2019). Adopters typically provide surrogate care for the orphan in the form of grooming, limited provisioning through food sharing, agonistic support, and even carrying (Samuni et al. 2019). As a result, adoption increases the survival probability of the orphan (Hobaiter et al. 2014). It is possible that the adoption of immature orphan chimpanzees also alleviates some of the effects of maternal loss on cortisol as observed in humans (Gunnar et al. 2001). Unfortunately, data were insufficient to evaluate effectively the effect of adoption on cortisol excretion profiles in this sample (adoption status is available for 9 out of 17 orphans, Table S1). Most orphans for whom we have data were adopted at some point during their immature years (Table S1) but the level of care provided by the adopter varies greatly across orphans (Samuni et al. 2019). Variation in the degree of care received by the orphans could explain the very large variation in cortisol levels observed between individuals within the first year after their mother’s death (**Figure S1**) but we do not have enough orphans in our study to assess the effect of the intensity of alloparental care.

In conclusion, our study provides evidence of an effect of early life adversity on cortisol levels and diurnal cortisol slopes in immatures of a wild mammal population. Interestingly, our study provides contrasts to studies on humans by showing apparent recovery from the impact of maternal loss on cortisol secretion profiles. This suggests that the biological embedding model proposed for humans may not apply to certain wild-living long-lived species, that prolonged alteration of the HPA axis functioning may not be viable in these species (Boonstra 2012; Dantzer et al. 2016; Beehner & Bergman 2017), and/or that access to buffering mechanisms alleviate the effect of early life adversity on cortisol profiles. In humans, orphans who did not have access to buffering mechanisms during childhood (such as placement in foster care family) present altered cortisol profiles as adults (Gunnar et al. 2001). Survival into adulthood for these human orphans, despite their altered physiology, may be enabled by access to societal systems such as medical care. Such systems are absent in non-human animals. The absence of altered cortisol profiles in adult orphan chimpanzees (our study) and baboons (Rosenbaum et al. 2020) suggest that the selective pressure for these orphans, and possibly for orphans in other wild long-lived non-human animal species, to return to regular HPA axis functioning, via buffering or other mechanisms, may be stronger than in humans.

There is much interest in linking proximate physiological responses to early life adversity to ultimate long-term fitness consequences (Dantzer et al. 2016). Whether and how the effect of maternal loss on immature, but not mature, cortisol profiles in wild chimpanzees contributes to the long-lasting negative impacts on their fitness (Walker et al. 2018; Crockford et al. 2020) remains to be established. In general, future studies on wild mammals should link the effects of early life adversity on an individual’s physiology to long-term fitness consequences to understand better the selection forces at play. An investigation of the physiological and social differences, between orphans who do survive and those who do not reach maturity, as well as identifying and quantifying the effects of the buffering mechanisms that contribute to these differences, will be key in this process.

## Material and Methods

### Study communities

We used the long-term data of the Taï Chimpanzee Project (Wittig & Boesch 2019b) collected on four communities of wild Western chimpanzees (East, North, Middle, and South) in the Taï National Park, Cote d’Ivoire (5°52′N, 7°20′E). The behavioural observation of the chimpanzees started in 1982 and is still ongoing. The observation periods for each of the communities are as follows: North 1982-present; South, 1993-present; Middle, 1995-2004; East, 2000-present (Wittig 2018). Urine samples were collected regularly in all communities from 2000 onwards except for the East community where sample collection started in 2003.

### Study subjects

For this study, we considered all immature individuals from both sexes (< 12 years of age) and mature males (> = 12 years of age) from whom urine samples were collected. The age range for immatures sampled in this study were 2.82-11.99 years for non-orphans and 4.10-11.99 years for orphans. Physical maturity may come later in male Taï chimpanzees but 12 years is the age at which chimpanzees range predominantly independently of their mother (Taï Chimpanzee Project, unpublished data) and are fully integrated in the male hierarchy (Mielke et al. 2018. We excluded mature females from the analysis since most females in our study immigrated from unhabituated communities into the study communities, which meant we had no knowledge of the presence or absence of their mothers during their immature years. We excluded orphans for whom the date of death of the mother was unknown (e.g. occurred before habituation of the study community). For individual samples, we excluded outliers (i.e. samples with very low or very high hormonal measures, see details in supplementary material) and samples collected when the individuals were sick. We also ensured that the final data set comprised at least three data points per individual per year and that the earliest and the latest samples were separated by at least 6 hours, to ensure a meaningful evaluation of the diurnal cortisol slope of each individual (see details in supplementary material). In total, we used 849 samples from 50 immatures, including 17 orphans (N samples per individual mean ± se = 17.0±2.2) and 2184 samples from 28 mature males, including 11 orphans (N samples per individual mean ± se = 78 ± 13.5).

### Demographic and behavioural data collection

In Taï each chimpanzee community is followed daily by a joint effort of local and international assistants and researchers (Wittig & Boesch 2019b). Each day, the observers conducted focal follows from and to sleeping sites. The focal individual was either followed all day (i.e. 12 hours) or the identity of the focal changed around 12h30 and two different individuals were followed each day one after the other (i.e. 6h focal follow each). The observer recorded detailed focal and *ad-libitum* behavioural data (Altmann 1974). Observers recorded all social interactions such as aggression and submissive behaviours, which we then used to build a dominance hierarchy (see below). In addition, each day, the observers recorded, the presence of all individual chimpanzees they encountered, which provides a detailed account of the demography of each community. Specifically, we obtained detailed information on individuals’ date of birth, immigration/emigration, and death or disappearance. This information was used to determine the early life history of the study subject, namely if their mother died before they reached 12 years of age, and, if so, the age of the subject when its mother died.

### Assessment of dominance hierarchy in mature males

Since dominance rank may correlate with the cortisol levels of adult male chimpanzees (e.g. Muller & Wrangham 2004) we wanted to control for this parameter in our analysis of cortisol patterns in adult males. We calculated the dominance hierarchy for mature males in each of the study communities using a modified version of the Elo-rating method (Neumann et al. 2011) developed by Foester et al. (2016). In this modified version, the *k* parameters and the starting score of each individual are optimised using maximum likelihood approximation (Mielke et al. 2018, see details in supplementary material). We used all of the long-term data available on unidirectional submissive pant-grunt vocalizations, given by the lower ranking of the two individuals towards the higher ranking (Bygott 1979). We used 9,189 pant-grunt recorded for males in Taï South, 3,952 in Taï East, 5,784 in Taï North, and 111 in Taï Middle. All Elo-rating scores were standardized between 0 and 1 with 1 being the highest-ranking individual and 0 the lowest ranking on any given day. We then extracted the Elo-rating score of each individual on the day when each urine sample was collected.

### Urine sample collection and analysis

During chimpanzee follows, we collected urine samples opportunistically from known individuals. Directly after urination, we collected the urine from leaves and/or the ground into a 2ml cryo vial using a disposable plastic pipette. Within 12 hours of collection, we placed these vials in liquid nitrogen. Subsequently, the samples were shipped on dry ice to the Endocrinology Laboratory of the Max Plank Institute for Evolutionary Anthropology in Leipzig, Germany, and stored at −80 °C until analysis. We used liquid chromatography mass spectrometry (LCMS, Hauser et al. 2008; Murtagh et al. 2013) and MassLynx (version 4.1; QuanLynx-Software) to quantify cortisol concentrations in each sample. For all samples analysed before September 2016, we corrected cortisol levels using prednisolone (hereafter the “old” method). Between September 2016 and June 2019, we corrected cortisol levels using cortisol d4 (hereafter the “new” method). To adjust for water content in the urine (i.e. urine concentration) we measured, for each sample, its specific gravity (SG) using a refractometer (TEC, Ober-Ramstadt, Germany). We corrected our cortisol concentration for urine water content in each sample using the following formula provided by Miller et al. 2004:

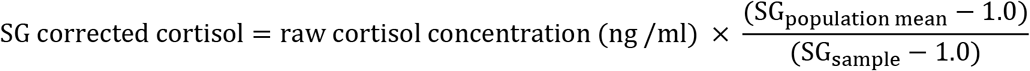

where SG_population mean_ is the mean SG value average across all the samples used in this study and SG_sample_ is the SG value of each given sample. In the manuscript, all cortisol concentrations are reported as ng/ml SG.

### Statistical analysis

We used a series of Linear Mixed Models (LMMs) to test our predictions regarding the effect of maternal loss on chimpanzee overall cortisol levels and diurnal slopes separately in immature males and females (Model 1a and 1b) and mature males (Model 2a and 2b). In all the models, each urine sample represented a data point and the cortisol concentration of the sample (expressed in ng/ml SG) was the response variable. We log-transformed the cortisol values to achieve a symmetric distribution of the response.

### Effect of maternal loss on cortisol profiles in immatures

In the first “immature” model (Model 1a), we tested the prediction that immature orphans will have higher overall cortisol levels and a flatter slope of diurnal cortisol slope when compared to non-orphans. We fitted a LMM with the orphan status (i.e. “no” if the mother of the individual was still alive on the day when the sample was collected and “yes” if the mother had died) as our test predictor to test for the effect of maternal loss on overall cortisol levels. In addition, to test for the effect of orphan status on the diurnal cortisol variation, we incorporated the linear and quadratic terms for time of sample collection as test predictors. Both the linear and quadratic terms for time of sample collection were included in interaction with orphan status into the model to test whether the diurnal cortisol slope differed between orphans and non-orphans. The time of sample collection was expressed in minutes with 0 being midnight and 720 being noon. In addition, in this model, we used the following control predictors: sex of the individual, community size, sex ratio of mature individuals in the community, age of the individual since this can influence the cortisol levels of immature chimpanzees (Tkaczynski et al. 2020), the LCMS method used (“old” or “new” method, see the *urine analysis* section). Community ID was not included as a control predictor in our analysis since it was highly correlated with community size. In addition, we accounted for seasonal variation in ecological conditions (e.g. rainfall, temperature, food availability) which can affect cortisol levels in chimpanzees (Wessling et al. 2018; Preis et al. 2019; Samuni et al. 2019) by converting the Julian date at which samples were collected into a circular variable and including the sine and cosine of this variable in our model (Wessling et al. 2018).

### Effect of age at mother’s death and time since mother died on cortisol profile in orphan immatures

In the second “immature model” (Model 1b), we focused on individuals who were orphans at the time of sampling in order to investigate more specifically the effects of the age at maternal loss, and years since maternal loss on cortisol levels. We tested the predictions that a) immature individuals who were orphaned at a younger age would have higher cortisol levels and flatter diurnal cortisol slopes and b) that, if some form of recovery occurs, these effects will be weaker the longer time has passed since an individual lost its mother. We incorporated two test predictors in Model 1b, the age at which the individuals have been orphaned (in days since their date of birth) and the time since the individuals have been orphaned (in days since the date their mother died). As in Model 1a, we incorporated the four interaction terms between these two test predictors and the linear and quadratic terms for time of sample collection. As before, we also incorporated individual sex, community size, sex ratio, and LCMS method, and the sine and cosine of the Julian date as control fixed effect. Initially, we also wanted to incorporate the age of the individual at the time of sampling into Model 1b. This was however not possible due to collinearity between the individual age at sample and both the age at which the individual was orphaned and the time since its mother died (i.e. the model did not run due to collinearity issues). To ensure that the results of Model 1b were not driven by the age at which the individual was sampled we ran an additional model (Model 1c) to investigate the relationship between individual age and cortisol levels and diurnal cortisol variation in non-orphan immatures. By doing so we could determine whether age-related changes in cortisol levels are expected to account for the patterns observed in Model 1b. In Model 1c, the cortisol level of each sample was the response variable and we used the age of the individual when the sample was collected as test predictor. As previously, we incorporated two interaction terms in the model between age and the linear and quadratic terms for time of day to test for the effect of age on cortisol levels and diurnal cortisol slopes. In addition, we also incorporated in Model 1c individual sex, community size, sex ratio, and LCMS method and the sine and cosine of the Julian date (i.e. seasonality) as control fixed effect.

### Effect of maternal loss on cortisol profiles in mature males

In the first “mature male” model (Model 2a), we tested whether mature males (> = 12 years) who were orphaned as immatures (i.e. before 12 years of age) had overall higher cortisol levels and a flatter diurnal cortisol slope than mature males who did not lose their mother before 12 years of age. We fitted a LMM with the early life orphan status (i.e. “no”, if the mother of the individual was still alive when the individual reached 12 years of age, and “yes” if the mother died before the individual was 12 years of age) as our test predictor. As in Model 1a, we incorporated two interaction terms between orphan status and the linear and quadratic terms for time of sample collection. As in the other models, we used community size, the sex ratio, the age of the individual, the LCMS method, and the sine and cosine of the Julian date as control fixed factors. In addition, we controlled for the dominance rank of the individual by adding the standardised Elo-rating score of each individual on the day the sample was collected as a fixed factor into the model.

### Effect of age at mother’s death on cortisol profile in orphan immatures

In the second “mature male” model (Model 2b) we assessed whether the age at which the orphan male chimpanzees lost their mother impacted the diurnal cortisol levels and slopes of mature males (i.e. whether the potential effect of early life adversity continued into adulthood). Accordingly, we fitted a LMM with “age at which mother died” as a test predictor and its interaction with the linear and quadratic terms for time of day. In this model, we used only samples collected from mature males who lost their mothers before they were 12 years of age. As previously, we used community size, the sex ratio, the age of the individual, the LCMS method, and the sine and cosine of the Julian date as control fixed effects.

In addition to the fixed effects, in all of the LMMs we included individual identity as a random factor to avoid pseudoreplication. To control for the changes in cortisol diurnal slope with age, we built one slope per individual per year into each model by incorporating as random factor a dummy variable “individual_year”. In addition, since certain years might have particularly harsh or favourable ecological conditions, and since this can affect cortisol levels in primates (e.g. Young et al. 2019), we also included year as a random factor in each model. Finally, our hormonal dataset included samples collected by different observers with different research interests (hereafter project). Thus, to account for potential variation in cortisol levels that may be a result of interobserver variation in the focus of research, we added the ‘project’ type as an additional random factor.

All analyses were conducted in R 3.6.2 (R Core Team 2018) using the function *lmer* from the package “lme4” (Bates et al. 2015). In each model, we included the maximal random slope structure between each fixed predictor (test and control) and each random effect (Baayen et al. 2008; Barr et al. 2013) but not the correlation between intercept and slopes. In particular, the linear and quadratic terms for time of day were included as random slopes within each of the random effects. In each model, we tested for the overall significance of the test predictors by comparing the full model to a null model comprising all control predictors, all the random effects, and random slopes but without the test predictors. We tested each full model against its corresponding null model using a Likelihood Ratio Test (LRT, Dobson 2002). We then assessed the significance of each predictor variable using a LRT between the full model and a reduced model comprising all the variables except the one to evaluate. This process was repeated across all variables, one by one, using the *drop1* function of the “lme4” package. If the LRT revealed that one interaction had a p-value > 0.05, we reran the model without this interaction and reassessed the significance of all the predictors.

Before fitting each model, we tested for collinearity issues between our predictor variables by computing the variance inflation factor (VIF) using the function *vif* from the package “car” (Fox & Weisberg 2011). Collinearity was not an issue in any of the final models (VIF of all predictor variables < 3.6). We also assessed model stability by removing one level of each random effect at a time and recalculating the estimates of the different predictors which revealed that the results were stable. For each model, we calculated the marginal R^2^ (i.e. the variance explained by the fixed effects) and the conditional R^2^ (i.e. the variance explained by the entire model including both fixed and random effects) using the function *r.squaredGLMM* of the package “MuMin” (Barton 2020).

## Acknowledgments

We thank the Ministère de l’Enseignement Supérieur et de la Recherche Scientifique and the Ministère de Eaux et Fôrets in Côte d’Ivoire, and the Office Ivoirien des Parcs et Réserves for permitting the study. We are grateful to the Centre Suisse de Recherches Scientifiques en Côte d’Ivoire and the staff members of the Taï Chimpanzee Project for their support and collecting the data. This study was funded by the Max Planck Society and the European Research Council (ERC) under the European Union’s Horizon 2020 research and innovation program awarded to C.C. (grant agreement no. 679787). Core funding for the Taï Chimpanzee Project has been provided by the Max Planck Society since 1997.

## Supplementary material

### Data preparation

The initial dataset comprised 4,604 samples (1,518 from immature individuals and 3,086 from mature males) from 46 female and 48 male immature individuals and 34 mature males. We applied a suite of selection criteria to subset our dataset to samples collected from individuals from whom all demographic and social data needed were available. We excluded all individuals for whom we could not assess if the mother died before they were 12 years of age or the age they were when their mother died. We also excluded samples for whom the cortisol concentration could not be measured or was excessively low (<0.1 ng/ml SG). We excluded samples with very low specific gravity (SG<1.003). Very low SG values are a sign of over diluted samples which reflect potential contamination with rain water and can, in turn, inflate cortisol concentration measurements. We also excluded samples collected from individuals on days when they displayed injuries or symptoms of sickness (as assessed by the on-site veterinary staff) since injury and sickness lead to extremely elevated cortisol levels in primates (e.g. Barton 1987; Muehlenbein & Watts 2010; Behringer et al. 2020). Finally, since a large part of our analysis focused on circadian cortisol variation, we excluded all samples for which we did not have a precise time of collection recorded. For the same reason, we limited our dataset for each individual to years when at least 3 samples were collected from this specific individual, and in years in which the earliest and the latest sample collections were separated in time by at least 6 hours. This criterion was applied in order to be able to calculate, in our statistical model, a meaningful circadian slope for each individual each year with time variation representing at least half of the active time of the chimpanzees (i.e. at least 6h out of 12h). The three samples could have been collected on different days but “time of sample collection” was used to define the 6 hour criteria. Following this selection process, we were left with 849 samples from 50 immatures (including 17 orphans) and 2184 samples from 28 mature males (including 11 orphans).

**Table S1.**
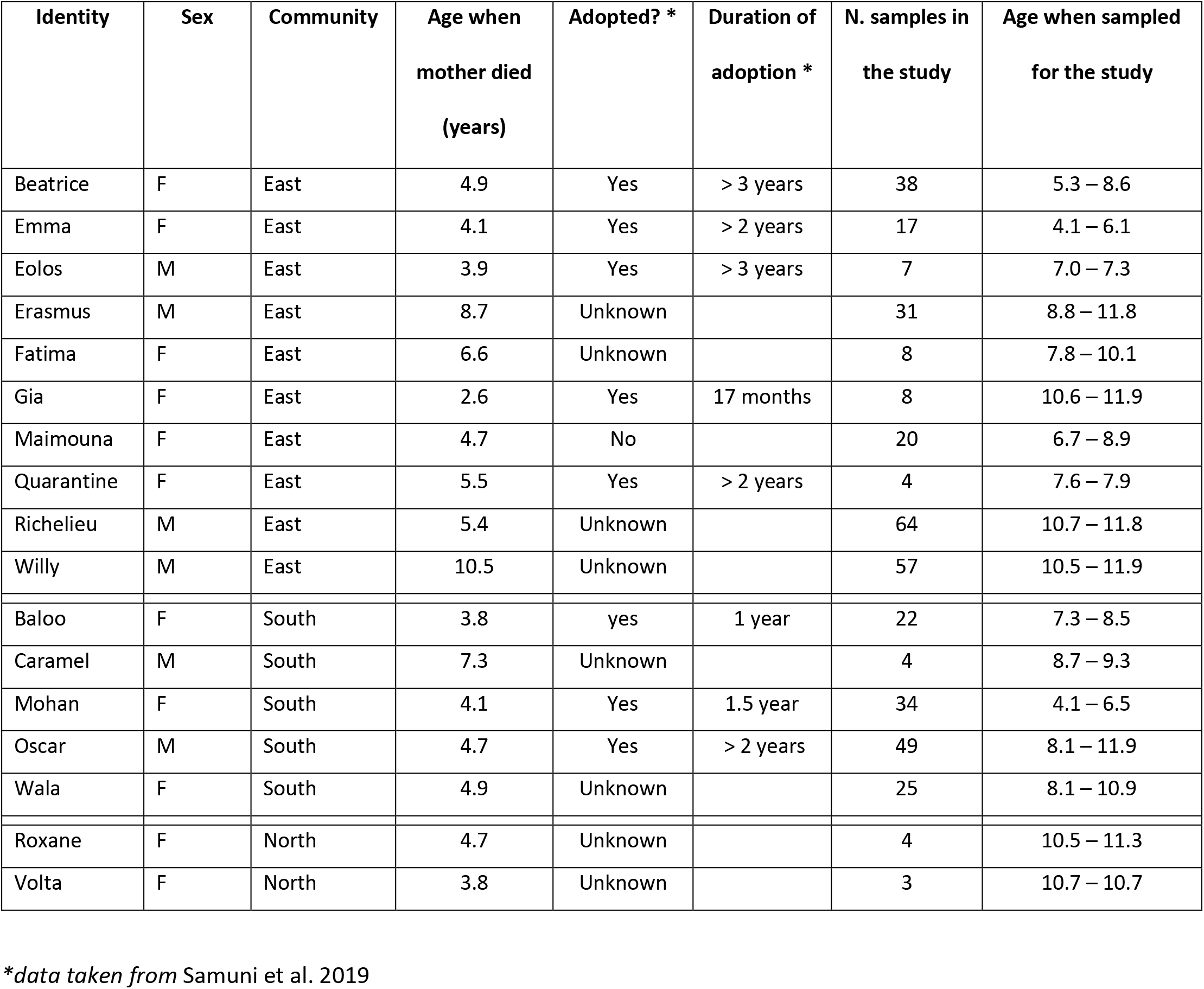
List of orphan immatures in the study and information about the adoption by adult individuals in the community.

| Identity | Sex | Community | Age when mother died (years) | Adopted? * | Duration of adoption * | N. samples in the study | Age when sampled for the study |
| --- | --- | --- | --- | --- | --- | --- | --- |
| Beatrice | F | East | 4.9 | Yes | > 3 years | 38 | 5.3 – 8.6 |
| Emma | F | East | 4.1 | Yes | > 2 years | 17 | 4.1 – 6.1 |
| Eolos | M | East | 3.9 | Yes | > 3 years | 7 | 7.0 – 7.3 |
| Erasmus | M | East | 8.7 | Unknown |  | 31 | 8.8 – 11.8 |
| Fatima | F | East | 6.6 | Unknown |  | 8 | 7.8 – 10.1 |
| Gia | F | East | 2.6 | Yes | 17 months | 8 | 10.6 – 11.9 |
| Maimouna | F | East | 4.7 | No |  | 20 | 6.7 – 8.9 |
| Quarantine | F | East | 5.5 | Yes | > 2 years | 4 | 7.6 – 7.9 |
| Richelieu | M | East | 5.4 | Unknown |  | 64 | 10.7 – 11.8 |
| Willy | M | East | 10.5 | Unknown |  | 57 | 10.5 – 11.9 |
| Baloo | F | South | 3.8 | yes | 1 year | 22 | 7.3 – 8.5 |
| Caramel | M | South | 7.3 | Unknown |  | 4 | 8.7 – 9.3 |
| Mohan | F | South | 4.1 | Yes | 1.5 year | 34 | 4.1 – 6.5 |
| Oscar | M | South | 4.7 | Yes | > 2 years | 49 | 8.1 – 11.9 |
| Wala | F | South | 4.9 | Unknown |  | 25 | 8.1 – 10.9 |
| Roxane | F | North | 4.7 | Unknown |  | 4 | 10.5 – 11.3 |
| Volta | F | North | 3.8 | Unknown |  | 3 | 10.7 – 10.7 |

*\*data taken from Samuni et al. 2019*

